# Real-world expectations and their affective value modulate object processing

**DOI:** 10.1101/408542

**Authors:** Laurent Caplette, Frédéric Gosselin, Martial Mermillod, et Bruno Wicker

## Abstract

It is well known that expectations influence how we perceive the world. Yet the neural mechanisms underlying this process remain unclear. Studies about the effects of prior expectations have focused so far on artificial contingencies between simple neutral cues and events. Real-world expectations are however often generated from complex associations between contexts and objects learned over a lifetime. Additionally, these expectations may contain some affective value and recent proposals present conflicting hypotheses about the mechanisms underlying affect in predictions. In this study, we used fMRI to investigate how object processing is influenced by realistic context-based expectations, and how affect impacts these expectations. First, we show that the precuneus, the inferotemporal cortex and the frontal cortex are more active during object recognition when expectations have been elicited a priori, irrespectively of their validity or their affective intensity. This result supports previous hypotheses according to which these brain areas integrate contextual expectations with object sensory information. Notably, these brain areas are different from those responsible for simultaneous context-object interactions, dissociating the two processes. Then, we show that early visual areas, on the contrary, are more active during object recognition when no prior expectation has been elicited by a context. Lastly, BOLD activity was shown to be enhanced in early visual areas when objects are less expected, but only when contexts are neutral; the reverse effect is observed when contexts are affective. This result supports the proposal that affect modulates the weighting of sensory information during predictions. Together, our results help elucidate the neural mechanisms of real-world expectations.

---

We expect to find hairdryers in bathrooms, tombstones in cemeteries, and baguettes in bakeries, but not tombstones in bathrooms, refrigerators in cemeteries and hairdryers in bakeries. That is, we live in a world where most objects are associated with specific contexts. Throughout a lifetime of experiences, we come to learn these associations, which lead us to form expectations about the objects to be encountered when we navigate the world. Congruent contexts have been shown to facilitate an object’s recognition, compared to incongruent contexts (Bar and Ullman, 1996; Biederman et al., 1982; Davenport and Potter, 2004); in addition, objects are recognized more accurately when a semantically consistent scene has been shown prior to the object’s recognition (Palmer, 1975; see Bar, 2004, for a review of related studies).

Perception can be understood as the process of integrating such top-down expectations with incoming sensory information. It has been proposed that predictions from high-level areas are transmitted to adjacent lower-level areas and compared with incoming sensory signals, such that only the discrepancy between these two signals – the prediction error – is transmitted up the visual hierarchy (Friston, 2005; see also Mumford, 1992; Ullman, 1995; Rao and Ballard, 1999). In support of this model, the expectation of a visual stimulus elicits a specific pattern of activity in the primary visual cortex (Kok et al., 2014, 2017; Hindy et al., 2016) and the perception of an expected stimulus results in reduced neural activity in sensory cortices (Summerfield et al., 2008; den Ouden et al., 2010; Alink et al., 2010; Kok et al., 2012a; Todorovic and de Lange, 2012; see de Lange et al., 2018, for a review). Some predictions, however, may require a different mechanism than feedback from adjacent visual areas (Bar, 2007; Hindy et al., 2016): for instance, the hippocampus has been shown to play a role in the generation of predictions (Hindy et al., 2016; Kok and Turk-Browne, 2018), and there is some evidence that parahippocampal (PHC) and retrosplenial (RSC) cortices initiate context-based expectations (Bar, 2003, 2004; Bar and Aminoff, 2003; Bar et al., 2006; Livne and Bar, 2016; Brandman and Peelen, 2017).

Most neuroimaging studies examining the mechanisms underlying predictions in visual perception have used very simple cues such as tones (Summerfield and Koechlin, 2008; den Ouden et al., 2010; Kok et al., 2012a, 2017) or a repetition of the same object (Summerfield et al., 2008; Todorovic and de Lange, 2012). By contrast, expectations about everyday objects usually stem from the surrounding context. Several previous studies investigated the processing of context-object relationships but did so by using a simultaneous presentation of the object and the scene (Faivre et al., 2019; Goh et al., 2004; Jenkins et al., 2010; Kirk, 2008; Rémy et al., 2014; see also Gronau et al., 2008, who use semantically related objects analyzed as a single event), which makes it hard to disentangle context-object interactions (occurring while both are being recognized) from context-based predictions (occurring prior to the object’s recognition). For example, context-object interactions can comprise matching processes or low-level crowding. While studying these processes is of interest, we focus here on predictions arising from an earlier processing of scenes and influencing a later processing of objects. To isolate this process, we used a sequential design in which scenes alone are presented before objects alone, just like when someone walks towards a cemetery and sees tombstones when he gets in. In fact, contexts are typically processed before the objects that they comprise (and they guide the eyes to these objects; Bar, 2003; Oliva & Torralba, 2007). Ganis & Kutas (2003) used a quasi-sequential scene-object presentation in their electroencephalography (EEG) study but their objects were shown overlayed to the scene previously presented; therefore, the effect of predictions on object recognition was not as isolated as in our design in which objects appear alone. They found a modulation of an N400-like component by the semantic congruency of the scene and object. Their use of EEG, however, prevented them from looking at the specific brain regions at play. To our knowledge, our study is the first neuroimaging study to explore context-based predictions using a sequential context-object design.

Relatedly, the effect of predictions has not been considered in the setting of an ecological object recognition task. Simple detection tasks (Jiang et al., 2013), delayed discrimination tasks (Kok et al., 2012a, 2014, 2017) or categorization tasks using few stimuli (den Ouden et al., 2010; Kok et al., 2012b) are typically used. Moreover, previous studies on prediction have manipulated predictability by artificial means, either by repeating and alternating stimuli (Summerfield et al., 2008; Todorovic and de Lange, 2012), by having stimuli appearing after different cues with different probabilities during the experiment (den Ouden et al., 2010; Kok et al., 2012a, 2012b, 2014, 2017; Jiang et al., 2013), or by developing arbitrary contingencies shortly before the experiment (Hindy et al., 2016). Associations between contexts and objects formed over a lifetime of experiences may involve mechanisms distinct from these. For instance, these associations learned a long time ago may be stored independently of the hippocampus and so, predictions would not originate from this area.

Moreover, real-world expectations are often tinted by some affective value. A visual context can elicit emotional reactions that may influence the recognition of objects in the scene (Lebrecht et al., 2012). In an emotional context (e.g., a cemetery), the affective value may be partially processed before the scene’s objects (e.g., a tombstone) and contribute to the object’s recognition (Barrett and Bar, 2009). In such a case, there would be a greater difference in brain activation between validly predicted and invalidly predicted objects for emotional contexts than for neutral contexts (due to the presence of additional affective information).

Alternatively, the prediction’s affective value might interact with its validity. Based on a recent proposal according to which sub-cortical nuclei have a modulating power over prediction errors (Kanai et al., 2015), and on the fact that many subcortical circuits are coordinated with bodily processes, Miller & Clark (2018) proposed that affect (relating to internal bodily states) exerts a continuous influence on perception by increasing the weight attributed to prediction errors during perceptual inference. When prediction errors are up-weighted, observers will rely more on the sensory input instead of their predictions to recognize a target object. This increased weight might be implemented by increasing the postsynaptic gain of neurons representing prediction errors (Kok et al., 2012b; Feldman & Friston, 2010). According to recent predictive coding models, the pattern of neural responses to valid and invalid predictions should reverse when prediction errors are given more weight (Feldman & Friston, 2010; Kok et al., 2012b). In our case, this would lead to a greater response to invalidly predicted stimuli than validly predicted stimuli when there is no affect, and to the opposite result when there is affect (see Kok et al., 2012b, for a description of a similar phenomenon, supplemented with diagrams).

In the present study, we aimed to address previous shortcomings by investigating how object recognition mechanisms are influenced by task-irrelevant implicit expectations generated by a predictive or non-predictive visual context. Which brain regions are responding to objects whether they were preceded by expectations or not will inform us about how high-level predictions are matched to sensory data. Furthermore, we aimed to investigate the effect of the affective value of predictions, and to disentangle between competing hypotheses on this matter. The pattern of results across the different affective value and predictive value conditions will teach us about how affect modulates the influence of predictions.

## Materials and Methods

### Participants

Seventeen healthy adults (9 female; mean age = 24.8; standard deviation = 4.3) were recruited on the campus of Aix-Marseille Université. Participants did not suffer from any neurological, psychological or psychiatric disorder and were free of medication. The experimental protocol was approved by the ethics board of CPP Sud-Méditerranée 1 and the study was carried in accordance with the approved guidelines. Written informed consent was obtained from all participants after the procedure had been fully explained, and a monetary compensation was provided upon completion of the experiment.

### Stimuli

We conducted several validation studies. For all these studies, subjects were French and had unlimited time to respond. In a first validation study, 35 different subjects (20 female; 22 to 54 years, mean = 29.2 years) were shown thirty-three context names and had to give the names of three objects with a high probability of being present in that context. Then, thirty-two public domain scene color images were selected from the internet as context images (see examples in Figure 1a). Context images were selected to ensure that their three most associated objects did not appear in them while still being representative of the context category.

**Figure 1.**
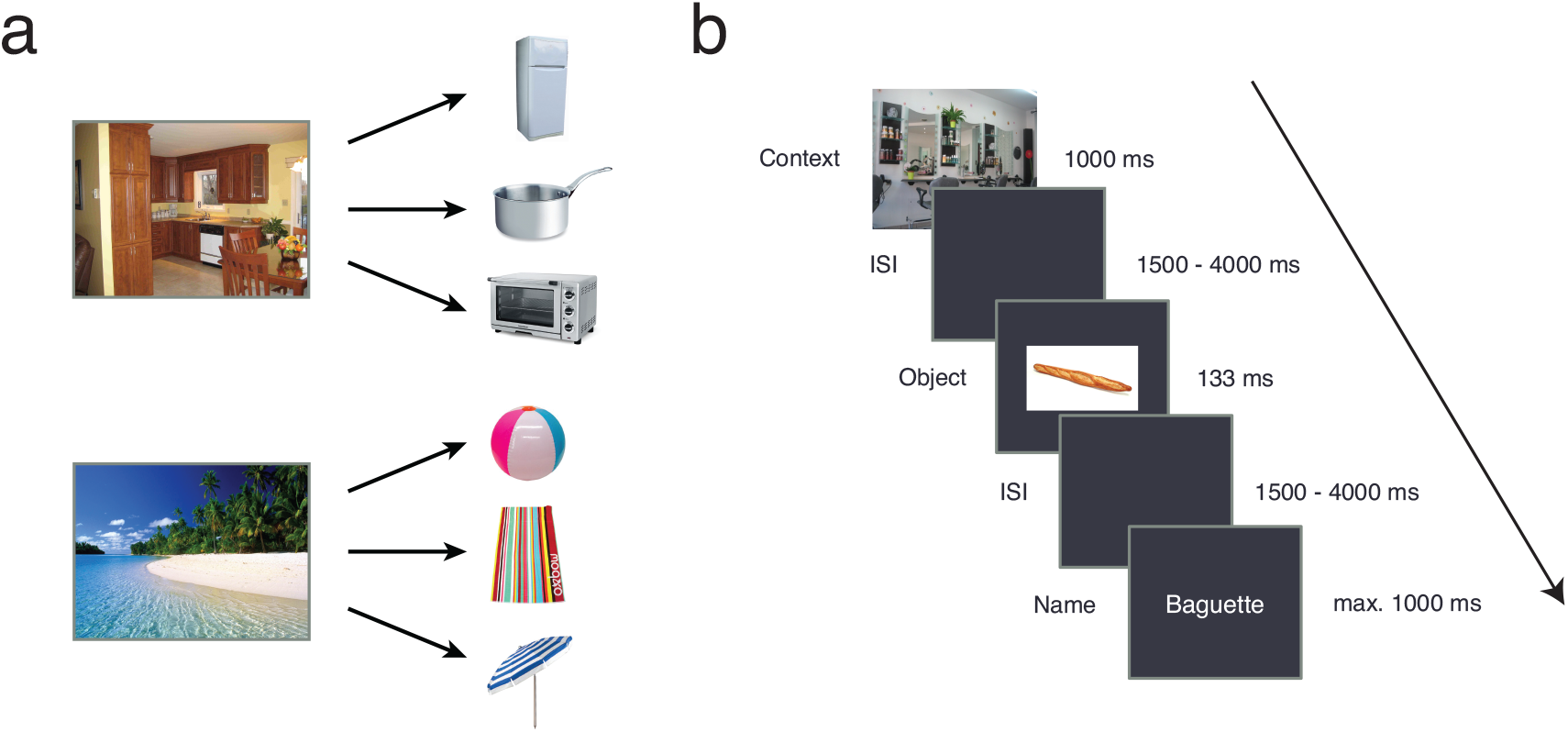
**A)** Example of one neutral and one affective scene, with their associated objects. See table S1 for a list of all contexts and associated objects. **B)** Example of a trial (Invalid Neutral condition). The object image and name have been enlarged for better viewing.

Ninety-six color images of objects corresponding to the three most cited names for each context were then selected for the experiment (e.g., swimsuit, diving board and pool ladder for swimming pool; Figure 1a, Table S1). For each context, the experimenters also chose three non-associated objects (selected from the ones that had never been associated with the context in the first validation study). Every object was the associated object of only one context and the non-associated object of only one other context; moreover, for each context, each one of the three non-associated objects was associated with a different context. Additionally, the experimenters (i.e. the four authors) divided contexts in Affective (*Aff*) and Neutral (*Neut*) categories according to their judgment of their affective intensity: the category of each context was determined by consensus.

We validated this categorization *a posteriori* with a second validation study. In this study, an independent sample of 22 subjects (12 female; 20 to 58 years, mean = 23.5 years) identified what they thought the context images represented (to confirm that the image represented the context; all contexts were largely correctly identified), indicated if the context elicited an emotion and, if so, what were its valence (negative to positive, from 0 to 10) and intensity (no emotion to very intense emotion, from 0 to 10); if the subject indicated that the context elicited no emotion, this was coded as 0 intensity. Valence was not included in the experimental design. We performed a Spearman correlation between our classification (binary variable) and the affective intensity score: we obtained a correlation coefficient of 0.78 solely with our binary classification, indicating that we explained a large part of the explainable variance; affective contexts had a mean intensity of 5.42 while neutral contexts had a mean intensity of 2.37. Looking more closely, only 4 contexts were miscategorized according purely to the validation’s intensity score; we repeated our fMRI analyses excluding these contexts and obtained very similar results (the regions activated were largely the same; see supplementary materials).

A third and final validation study was conducted to collect quantitative measures of the associations between objects and contexts. Forty-four new subjects (29 female; 18 to 86 years, mean = 40.7 years) indicated on a scale from 0 to 10 exactly how much each object was associated to its associated and non-associated contexts (context-object pairs were randomized). Measures were z-scored within each subject. Associated objects had a vastly greater associative weight with their context (F = 38353, *p* < 4.4 × 10^−65^). Neutral contexts had a greater association to their associated objects on average, and a smaller association to their non-associated objects (significant interaction between association and affective value; F = 17.43, *p* = .0001). To get rid of this potential confound, we used these weights, z-scored within predictive value condition but not within affective value condition, as covariates in the analysis.

Finally, we randomized the phases of the mean of the context images in the Fourier domain – separately for each RGB color channel – to obtain 96 different phase-scrambled images.

### Data acquisition

Functional imaging data were acquired with an AVANCE 3 Tesla scanner (Bruker Inc., Ettlingen, Germany) equipped with a 2-channel head-coil. Functional images sensitive to BOLD contrast were acquired with a T2*-weighted gradient echo EPI sequence (TR 2400 ms, TE 30 ms, matrix 64 × 64 mm, FOV 192 mm, flip angle 81.6°). Thirty-six slices with a slice gap of 0 mm were acquired within the TR; voxels were 3 × 3 × 3 mm. Between 303 and 311 volumes were acquired in each run, excluding the six dummy scans acquired at the beginning of each run for signal stabilization. Additionally, a high resolution (1 × 1 × 1 mm) structural scan was acquired from each participant with a T1-weighted MPRAGE sequence.

### Experimental Design

The LabVIEW (National Instruments Inc., Austin, TX, USA) software was used to project stimuli during the experiment. Stimuli were projected to a screen positioned in the back of the scanner using a video projector. Subjects could see the video reflected in a mirror (15 × 9 cm) suspended 10 cm in front of their face and subtending visual angles of 42 degrees horizontally and 32 degrees vertically.

Each trial was built as follows: a large cue image (see below) spanning the whole screen during 1 s, a blank screen during 1.5 to 4 s (duration randomly selected from a truncated exponential distribution with mean of 2 s), a centered object image on a white background (20 × 26 degrees of visual angle) during 133 ms, a blank screen during 1.5 to 4 s, and an object name shown until the subject answered or for a maximum of 1 s (Figure 1b). During the whole trial except at the presentation of the full-screen scene, a black background was present around the stimuli. Subjects answered by pressing one of two buttons on a hand-held response device (as accurately and rapidly as possible) to indicate if the name corresponded to the object, which occurred on 80% of the trials. The purpose of the task was to maintain the attention and engagement of the subjects during the perception of the scenes and objects. A black screen was displayed for an additional 1 s between trials.

On a third of the trials (*Valid* condition), the cue image was a scene associated with the object following it (e.g., an airport and a suitcase); on another third (*Invalid* condition), it was a scene not associated with the object following it (e.g., a church and a tennis racket); on the final third (No-Prediction condition; *noPred*), it was a scrambled image (always a different one). Each object was shown once in each of these conditions, for a total of 288 trials.

Trials were divided into three runs of 96 trials each. The order of the trials within each run was predetermined randomly prior to the study and was constant across participants; the order of the three runs was counterbalanced across subjects. Each functional run lasted between 10 and 12 mins (differences in length were due to the random inter-stimuli intervals), with short breaks between them.

### Data Preprocessing and Analysis

The SPM8 software (http://www.fil.ion.ucl.ac.uk/spm/), running in the MATLAB environment (Mathworks Inc., Natick, MA, USA), was used. T1-weighted structural images were segmented into white matter, gray matter and cerebrospinal fluid, and warped into MNI space. Functional images were realigned, unwarped and corrected for geometric distortions using the field map of each participant, slice time corrected, coregistered to the structural image of the corresponding participant, and smoothed using an isotropic Gaussian kernel with a 6 mm Full-Width Half-Maximum.

A standard General Linear Model (GLM) analysis was performed for each subject. Three events were modelled on each trial: contexts (or scrambled images), objects and object names. Object events (the regressors of interest) were modelled for each condition separately (Valid-Aff, Valid-Neut, Invalid-Aff, Invalid-Neut and noPred); scene events (regressors of no interest) were also modelled separately for each condition; one additional regressor was included for the object names. All these events were modelled as Dirac delta functions (duration of zero) convolved with SPM8’s canonical hemodynamic response function. To get rid of potential effects caused by differences in context-object associations, we included an additional parametric regressor which consisted of the context-object association weights as determined by our third validation study. This regressor was z-scored separately within Valid contexts and Invalid contexts but not separately within each affective value subcondition so that differences in context-object associations between affective and neutral contexts were accounted for, but that differences between Valid and Invalid conditions remained; finally, we convolved it with the hemodynamic response function. The six motion parameters were also included as additional nuisance regressors.

A temporal high-pass filter (cut-off of 128 s) was used to remove low-frequency drifts, and temporal autocorrelation across scans was modelled with an AR(1) process. Contrasts were then computed at the subject level and used for group analyses using one-sample *t*-tests. All voxels inside the brain were analyzed; we maintained the familywise error rate of *p* < .05, two-tailed, at the cluster level (primary threshold of *p* < .001, uncorrected) using random field theory (Friston et al., 1994). The Anatomy (Eickhoff et al., 2005) and WFU-PickAtlas (Maldjian et al., 2003) toolboxes were used to identify activated brain regions based on peak Montreal Neurological Institute (MNI) coordinates.

Data and code will be made available on the Open Science Framework repository shortly after the publication of this manuscript.

## Results

### Behavioral results

Mean accuracy was 97.1% (σ = 2.8%) for the Valid-Neut condition, 97.2% (σ = 2.1%) for the Valid-Aff condition, 96.9% (σ = 2.6%) for the Invalid-Neut condition, 95.0% (σ = 2.5%) for the Invalid-Aff condition and 95.7% (σ = 2.4%) for the noPred condition. When comparing Valid, Invalid and noPred together (ANOVA, *n* = 17), there was no effect of condition on accuracy (*F*(2,16) = 2.17, *p* = .26, *η*^*2*^_*p*_ = 0.12). When comparing all subconditions except noCont together in order to include affective value as a factor (ANOVA, *n* = 17, corrected for multiple comparisons across both ANOVAs), there was a significant main effect of predictive value (*F*(1,16) = 10.36, *p* = .01, *η*^*2*^_*p*_ = 0.13), i.e. accuracy was higher when objects were predicted. There was no significant main effect of affective value (*F*(1,16) = 4.42, *p* = .10, *η*^*2*^_*p*_ = 0.08), and no significant interaction between affective and predictive values (*F*(1,16) = 3.97, *p* = .13, *η*^*2*^_*p*_ = 0.10).

Mean response time was 640 ms (σ = 121 ms) for the Valid-Neut condition, 650 ms (σ = 120 ms) for the Valid-Aff condition, 638 ms (σ = 110 ms) for the Invalid-Neut condition, 636 ms (σ = 127 ms) for the Invalid-Aff condition and 632 ms (σ = 116 ms) for the noPred condition. When comparing Valid, Invalid and noPred (ANOVA, *n* = 17), there was no effect of condition on response time (*F*(2,16) = 1.53, *p* = .46, *η*^*2*^_*p*_ = 0.09). When comparing all conditions except noPred together together in order to include affective value as a factor (ANOVA, *n* = 17, corrected for multiple comparisons across both ANOVAs), there was no main effect of affective or predictive value and no interaction (*F*s(1,16) = 1.67, 0.25 and 0.64 respectively, *p* > .40, *η*^*2*^_*p*_ < .03).

### fMRI results

We first investigated which regions responded to the presentation of objects in general. All object presentations (irrespectively of the specific condition) were contrasted against the implicit baseline. Several areas were active, most notably the bilateral fusiform gyri and parts of the left occipital cortex (see supplementary materials). These areas are part of the lateral occipital complex, which is typically activated during object perception (Grill-Spector et al., 2001).

To investigate the potential effect of explicit contextual expectations, irrespectively of their validity, on brain activity during object recognition, we contrasted the Valid and Invalid conditions with the noPred condition (paired t-test, *n* = 17). This contrast reveals areas where expectations and sensory information are combined or integrated, i.e. areas where sensory processing is modulated by expectations irrespectively of their validity (Summerfield & Koechlin, 2008). Five clusters were significantly more activated in the Valid and Invalid conditions than in the noPred condition (*p* < .05, two-tailed, corrected for family-wise error rate (FWER); peak Cohen’s *d*_*z*_ = 1.91; Figure 2; Table 1): one bilateral cluster in the precuneus (which is part of the retrosplenial complex), one extending from the left precuneus and middle occipital gyrus to the left angular gyrus, one in the left middle temporal gyrus, one in the left middle and inferior frontal gyri and one in the right angular gyrus. The reverse contrast revealed the specific activation of two clusters in the right superior and middle occipital gyri and in the left middle occipital gyrus (*p* < .05, two-tailed, FWER-corrected; peak Cohen’s *d*_*z*_ = 1.73; Figure 2; Table 1). To confirm that our randomized jitters prevented contamination of object responses by scene information, we also conducted supplementary analyses comparing the same conditions but at the moment of context presentation instead of object presentation (i.e. contrast of scenes to scrambled scenes). We observed the activation of occipital areas and most notably, of the parahippocampal gyrus, a region typically associated to scene processing (Epstein et al., 2003; see supplementary materials). Importantly, these regions were only partly overlapping with those engaged at the moment of object presentation, and the scene-processing parahippocampal place area was only implicated when directly contrasting scenes.

**Figure 2.**
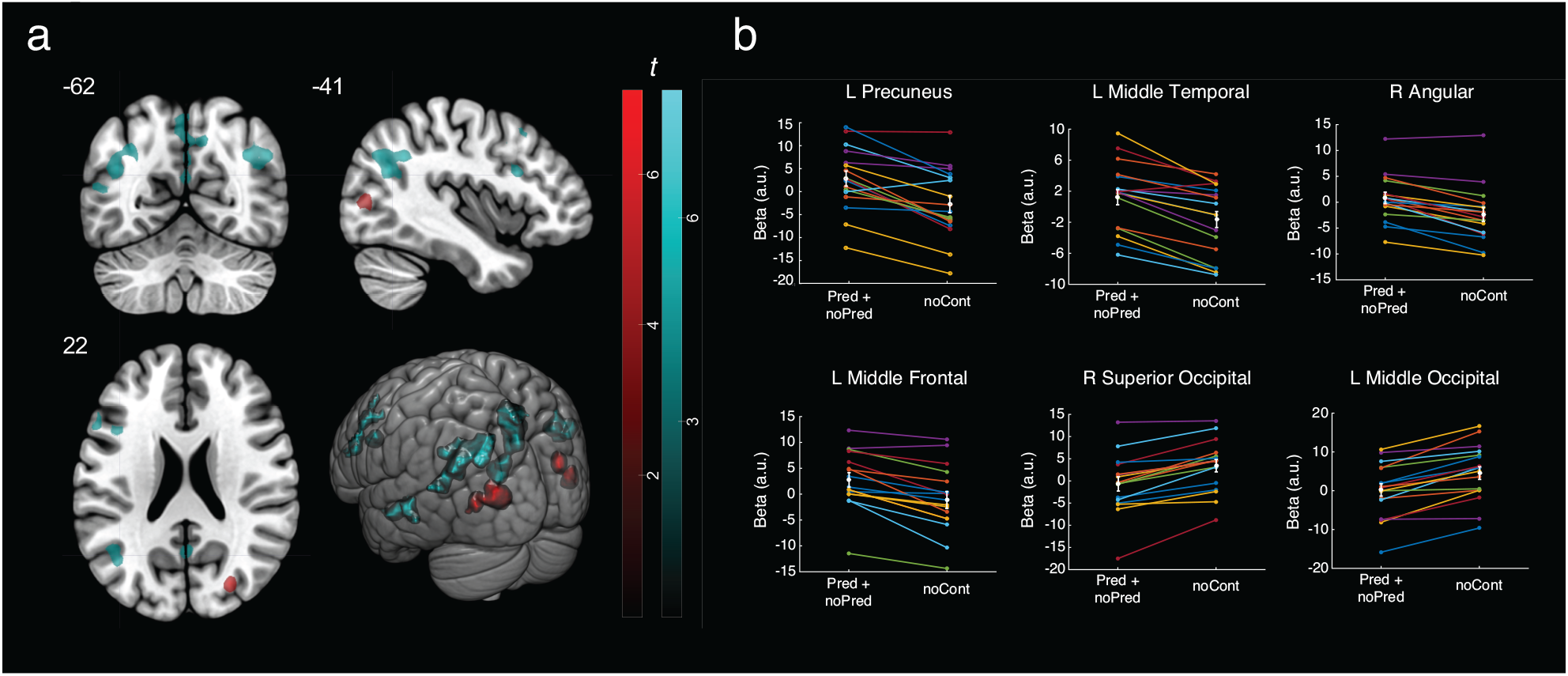
**A)** Significant clusters for the noCont > (Pred + noPred) (in red) and the (Pred + noPred) > noCont (in cyan) contrasts. **B)** Beta values of individual subjects for noCont and (Pred + noPred) conditions in peak voxels of various significant clusters, along with the group means and standard errors.

**Table 1.** Montreal Neurological Institute (MNI) coordinates and T values for significantly activated brain regions.

| Brain regions | Peak MNI coordinates |  |  | Nb of voxels | Peak T value |
| --- | --- | --- | --- | --- | --- |
|  | x | y | z |  |  |
| *(Pred + noPred) > noCont* |  |  |  |  |  |
| L Middle Occipital | -30 | -69 | 39 | 232 | 7.88 |
| L Angular | -42 | -63 | 27 |  | 6.63 |
| L Middle Temporal | -54 | -21 | -9 | 76 | 6.92 |
| R Angular | 51 | -63 | 36 | 90 | 6.16 |
| L Middle Frontal | -48 | 18 | 39 | 94 | 5.79 |
| L Inferior Frontal | -42 | 15 | 24 |  | 5.51 |
| L Precuneus | -3 | -66 | 54 | 224 | 5.59 |
| R Precuneus | 9 | -57 | 42 |  | 5.32 |
| *noCont > (Pred + noPred)* |  |  |  |  |  |
| R Superior Occipital | 27 | -78 | 21 | 61 | 7.12 |
| R Middle Occipital | 36 | -81 | 12 |  | 4.75 |
| L Middle Occipital | -27 | -84 | 6 | 90 | 6.76 |
| *Predict. x Affect.* |  |  |  |  |  |
| R Cuneus | 15 | -96 | 6 | 45 | 7.79 |
| L Cuneus | -12 | -90 | 3 | 43 | 6.08 |
|  | 0 | -87 | -3 |  | 4.55 |
|  | -9 | -84 | -9 |  | 3.89 |

We then investigated whether there was a main effect of predictive value (Valid vs Invalid), a main effect of the context’s affective value (Aff vs Neut), and an interaction between predictive and affective values at the moment of object recognition (paired t-tests, *n* = 17). There were no significant main effects of predictive or affective values. However, there was a significant interaction between predictive and affective values for two clusters: one in the right cuneus and one overlapping the left cuneus, calcarine gyrus and lingual gyrus (*p* < .05, two-tailed, FWER-corrected; peak Cohen’s *d*_*z*_ = 1.74; Figure 3; Table 1). We then investigated the simple effects specifically in the peak voxels (local maxima) of the clusters which had a significant interaction effect. We observed that these voxels were more active in the Valid-Aff condition than in the Invalid-Aff condition (left cuneus: *t*(16) = 4.80, *p*_*Bonf*_ = .0008, *d*_*z*_ = 1.16; right cuneus: *t*(16) = 4.56, *p*_*Bonf*_ = .001, *d*_*z*_ = 1.11) and more active in the Invalid-Neut than in the Valid-Neut condition (left cuneus: *t*(16) = 4.41, *p*_*Bonf*_ = .002, *d*_*z*_ = 1.07; right cuneus: *t*(16) = 4.24, *p*_*Bonf*_ = .003, *d*_*z*_ = 1.03).

**Figure 3.**
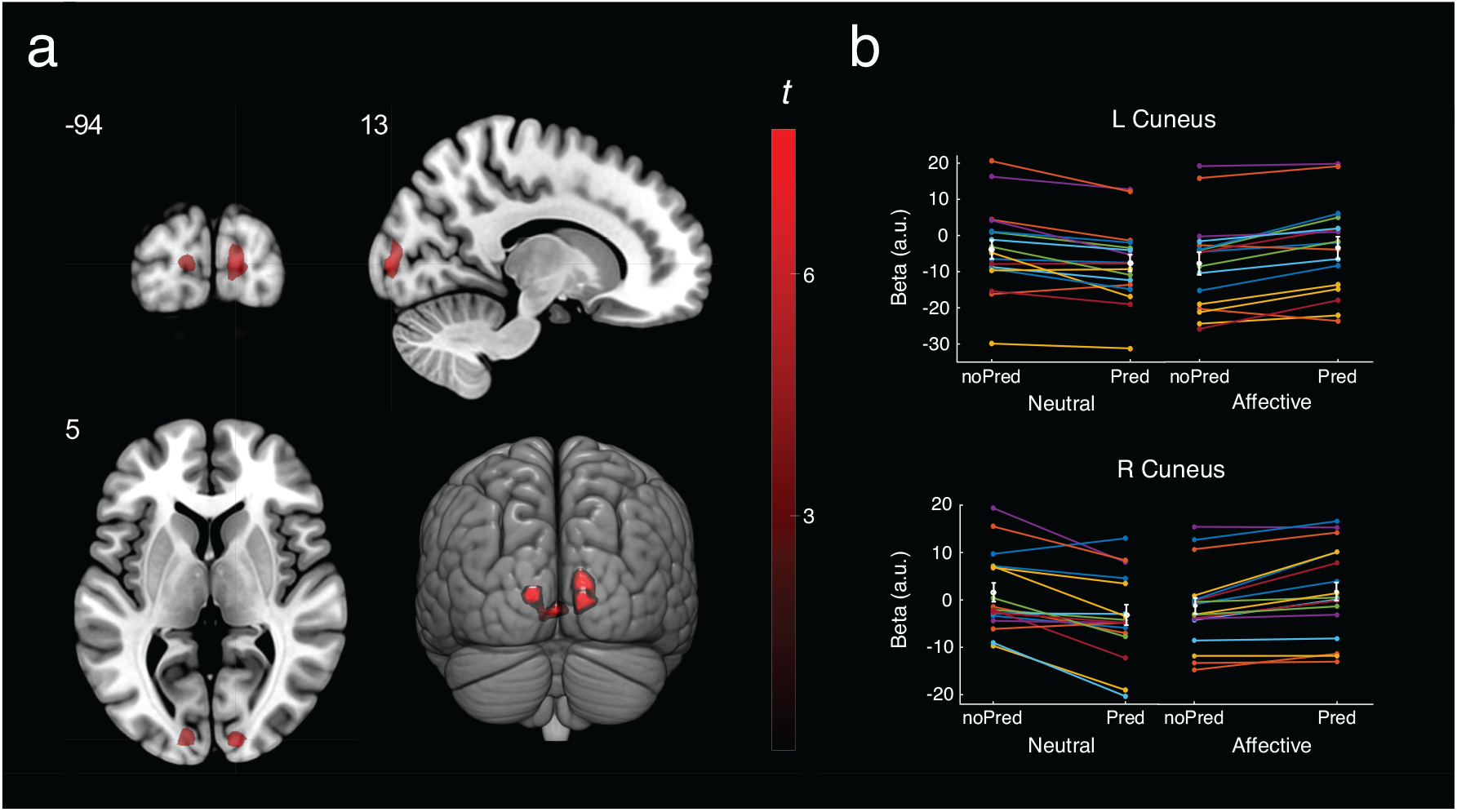
**A)** Significant clusters for the interaction between affective and predictive values (in red). **B)** Beta values of individual subjects for each condition in peak voxels of significant clusters, along with the group means and standard errors.

Next, we conducted a series of control analyses to ensure that the interaction could not have been the result of undesirable confounds. First, we investigated whether the interaction could have been caused by differences between the objects associated to neutral contexts and those associated to affective contexts by assessing if there was any significant difference in brain activity when they were perceived without a context (noPred condition). There was no significant difference between the conditions (*p*_FWER_ > .33). We also analyzed the image similarities directly: we used the HMAX model (Riesenhuber & Poggio, 1999; Serre et al., 2007), a commonly used model of the early visual cortex, and we computed correlation distances between the responses of the model to each image. We then verified if the between-categories (affective context objects to neutral context objects) distances were larger than the within-categories distances (two sample t-tests): no difference was observed (compared to within-neutral distances: .498 vs .504, *t*(3430) = .57, *n* = 1128 and 2304, *p* = .57; compared to within-affective distances: .498 vs .502, *t*(3430) = .38, *n* = 1128 and 2304, *p* = .70), suggesting that the images in these two object categories are similar.

Finally, the possibility remained that attention could explain the interaction between affective and predictive values: a similar interaction has indeed been previously reported with attention as a factor instead of affective value (Kok et al., 2012b). A first objection to this claim would be that our behavioral results actually trend (*p* =.10) toward an opposite effect. Although we observe the same reversed prediction effect in the brain for affective contexts that Kok et al. observed for task-relevant stimuli, the lower recognition accuracy in the affective condition suggests that they are not attended more and that attention is not the cause of this interaction. Nonetheless, we decided to conduct an additional behavioral experiment to isolate potential attentional effects better. Twenty-four participants aged between 18 and 30 years (mean = 25.2) performed a Gabor orientation discrimination task (vertical vs horizontal), in which the Gabor patches (1 cycle per degree) were randomly following either a neutral context image or an affective context image in the same way as in the fMRI experiment (contexts presented for 1s, 1.5-4 s jitter, patches presented during 133 ms). Subjects were instructed to respond as accurately and rapidly as possible, and adaptive procedures were conducted separately in each condition in order to find the contrast sensitivity threshold associated with each condition. Again, no difference was observed (log_10_(contrast) of −2.10 vs −2.11; *p* = .94). Since we know that contrast sensitivity is greatly enhanced by attention (see Carrasco, 2006, for a review), it does not seem likely that affective contexts were attracting attention and maintaining it for up to 4 seconds in order for it to alter object processing.

## Discussion

Our first aim was to investigate how the generation of expectations about objects from a preceding context might modulate the activity of brain areas involved in object perception. We found significantly more activation in the precuneus, the left middle occipital gyrus, the left middle temporal gyrus, the left frontal cortex and the parietal cortex, when (valid or invalid) contextual expectations were generated prior to object perception, suggesting that these high-level areas are mainly associated with object processing when expectations are generated. These activations specifically represent an interaction between contextual expectations and object bottom-up sensory information: activity related solely to object processing is cancelled out because the objects are the same in both conditions, and activity related solely to the prior presentation of the context is regressed out in the GLM.

To our knowledge, only Summerfield & Koechlin (2008) performed a similar analysis before; however, they used lines as cues and gratings as stimuli, and the cue was directly related to the task (the subjects had to indicate whether the cue and the grating matched). In their study, they observed a significantly greater activation of the middle occipital and fusiform gyri when there was an expectation. We also find a greater activation of the middle occipital gyrus, in addition to many other brain regions. Since expectations in our study are about objects rather than simple grating orientations, regions representing them are likely to be more numerous. The interaction between object and context processing observed in the middle temporal gyrus (a part of the inferotemporal cortex) supports a popular hypothesis according to which top-down contextual predictions would be combined with bottom-up sensory information to facilitate object recognition in the inferotemporal cortex (Bar, 2004). The precuneus and the parietal cortex, which are also activated in this contrast, have previously been linked to episodic memory retrieval and contextual associative processing (Lundstrom et al., 2005; Aminoff et al., 2007; Livne and Bar, 2016; Brandman and Peelen, 2017) which both require the integration of stored representations with incoming sensory information. Moreover, the precuneus of an observer that views several objects simultaneously is more activated when these objects are contextually related than when they are not (Livne and Bar, 2016); this suggests that the contextual representations elicited by some of these objects are compared to other objects. Recently, activity in the retrosplenial complex, a region comprising the precuneus, has been shown to correlate with supra-additive decoding of objects embedded in scenes, suggesting that the precuneus is responsible for a scene-based facilitation of object representations (Brandman and Peelen, 2017). Interestingly, the interaction we observed between context and object information in the precuneus is also supra-additive (i.e. there is a remaining positive activation after considering the main effects of object and context). We extend previous results by showing that the precuneus integrates object sensory information with valid or invalid scene-based expectations generated prior to object presentation. The inferior and middle frontal gyri were also active during object processing when expectations were generated. These regions have previously been found to respond more to objects in non-congruent scenes than to objects in congruent scenes (Rémy et al., 2014): it is thus likely that they are responsible of integrating contextual information with perceived objects. The prefrontal cortex has also been linked to object-related predictions in other studies (Bar, 2007; Bar et al., 2006) and has been found to both maintain expectations and integrate them with sensory information (Summerfield et al., 2006; Summerfield & Koechlin, 2008).

The reverse contrast, associated with visual processing of objects when no expectation (neither valid nor invalid) had been generated from a context, yielded bilateral activation of primary visual areas. Activated voxels may be part of areas primarily associated with the processing of sensory information shared by a majority of objects (e.g., intermediate spatial frequencies; Caplette et al., 2014), which is thus reduced when almost any object is expected.

We then investigated whether there was an effect of prediction error or match, i.e. whether some areas were more active at the presentation of the object when the object followed a valid context or when the object followed an invalid context. When neutral and affective contexts were combined, there was no significant difference between valid and invalid conditions; however, there was a significant interaction between predictive and affective values in low-level occipital areas, specifically the left and right cunei. Looking at this cluster, the classical prediction error effect was visible for neutral contexts, i.e. validly predicted objects elicited a smaller BOLD signal; but, when contexts were affective, this effect was reversed, i.e. validly predicted objects elicited a larger BOLD signal. Note that previous studies observing a smaller signal for validly predicted objects have exclusively used affectively neutral cues, making our results compatible with theirs.

These results are not compatible with the proposal that a subject’s internal affective state is altering the content of their predictions about object identities (Barrett and Bar, 2009). According to this idea, the affective value of a preceding context (or even a simultaneous context or the object itself; see Barrett and Bar, 2009) would alter the subject’s bodily state and bring additional information that could be used by the brain to predict the identity of perceived objects. Consequently, a similar pattern of results should be visible for neutral and emotional contexts, with only a greater difference in activation between validly predicted and invalidly predicted objects for emotional contexts than for neutral contexts (due to the additional emotional information).

Our results are compatible, however, with the general idea that affect interacts with predictive processing (Barrett and Simmons, 2015; Miller and Clark, 2018). One possibility recently put forward by some authors is that, rather than contributing to the content of the predictions, a subject’s internal affective state modulates the weight given to prediction errors during perceptual inference (Miller and Clark, 2018). Specifically, this weight would increase when the subject is experiencing affect. According to recent predictive coding models, such an increased weighting should lead to a reversal of the classical prediction error effect, as we observed (Feldman & Friston, 2010; Kok et al., 2012b).

Note that most affective contexts (11/15) used in our experiment had a positive valence. This may have affected the results. However, because negative contexts were especially negative, the median valence for affective contexts was still close to neutral (5.8, 0 being the most negative and 10 the most positive). Moreover, Miller & Clark (2018) do not differentiate between positive and negative stimuli in their proposal and, generally, the same brain network seems to be implicated in the processing of all affective stimuli, irrespective of valence (Lindquist et al., 2016).

Kok and colleagues (Kok et al., 2012b) reported a similar reversal of the prediction error effect in the early visual cortex for task-relevant stimuli. They argued that this effect was caused by endogenous attention increasing the weight of prediction errors (Friston, 2009; Feldman & Friston, 2010; Rao, 2005). This cannot be the cause of the effect we observed however, since attention was not manipulated in our study and our stimuli were all similarly task relevant. Furthermore, exogenous attention was also similar between our conditions, as revealed by behavioral results obtained in the scanner and in the control contrast sensitivity experiment. This implies that the reversal of the prediction error effect in our study was not caused by an increase in attention.

Notably, we observed significant main effects neither of predictive nor of affective value. The absence of a main effect of prediction can be explained by the fact that the effect of prediction is completely reversed depending on the affective value and so is cancelled when all contexts are combined. The absence of a main effect of affect might be due at least partly to the fact that our contrast concerns object processing and that affective value was at the level of the context. However, we also did not observe the activation of several regions typically implicated in affect processing during affective scene presentation (see supplementary materials). Our scenes, which were for the most part everyday contexts, possibly did not have a sufficient intensity to elicit such activations: the stimuli used in most fMRI studies are often of a much higher average intensity (e.g., IAPS; Lang et al., 2005).

In summary, real-world expectations initiated by contexts, irrespectively of their degree of validity, led to more activation of high-level areas (including parietal and frontal cortices) during subsequent object recognition; notably, these regions were distinct from those responsible of instantaneous context-object interactions. It is important however to note that these prior expectations are not necessarily explicit and conscious: they are representations preactivated by the perception of the scene, which will impact the processing of subsequent stimuli. In addition, the context’s affective value interacted with the validity of the prediction it had initiated: classical prediction error effects were only observed with neutral contexts, and a complete reversal of these effects was observed when contexts were emotional. This result is not compatible with the idea that the affective value of a stimulus, and the ensuing internal bodily state of the subject, are contributing to the creation of predictions (Barrett and Bar, 2009); but it is compatible with a modulatory role of affective value over the weight of sensory evidence in perception (Miller and Clark, 2018). In conclusion, our results deepen our understanding of predictive coding in an ecological setting by showing that the mere presence of explicit expectations, and their affective content, modulate object recognition.

## Acknowledgements

This study was funded by the Fondation Planiol (BW), by the Institut Universitaire de France (MM) and by the Social Sciences and Humanities Research Council of Canada (LC).

