## Supplemental Material for "Real-world expectations and their affective value modulate object processing"

### Supplementary Materials

**Table S1.** Objects associated to each context.

| *Affective Contexts* | **Intensity** | **Valence** | | **Associated objects** | |
|---|---|---|---|---|---|
| **Cemetery*** | 4.0 | 2.9 | Tombstone | Wreath | Memorial plaque |
| **Funfair** | 4.4 | 7.2 | Cotton candy | Bumper car | Ferris wheel |
| **Fire** | 7.1 | 1.6 | Fire truck | Fire hose | Fire extinguisher |
| **Luxury Hotel*** | 3.3 | 7.1 | Crystal chandelier | Large bed | Champagne bottle |
| **Ski Slope** | 6.2 | 7.8 | Skis | Chair lift | Ski poles |
| **Dumping ground** | 6.0 | 1.6 | Plastic bag | Old shoe | Garbage truck |
| **Circus** | 5.6 | 7.5 | Clown nose | Trapeze | Hoop |
| **Beach** | 7.2 | 8.8 | Beach towel | Parasol | Beach ball |
| **Nursery** | 6.1 | 8.2 | Teddy bear | Mobile | Changing mat |
| **Stage** | 6.1 | 7.6 | Electric guitar | Microphone | Speakers |
| **Wedding** | 5.1 | 7.3 | Wedding cake | Wedding dress | Wedding bouquet |
| **War zone** | 6.1 | 1.3 | Machine gun | Military helmet | Grenade |
| **Birthday** | 4.3 | 6.7 | Birthday cake | Present | Birthday candles |
| **Nightclub** | 4.4 | 6.8 | Disco light | Disco ball | Cocktail glass |
| **Hospital room** | 6.1 | 2.3 | Syringe | Stethoscope | Surgical mask |
| **Boat** | 4.9 | 7.4 | Life buoy | Rudder | Life vest |
| ***Neutral Contexts*** | | | | | |
| **Hairdresser** | 1.4 | 6.1 | Scissors | Comb | Hairdryer |
| **Office** | 1.9 | 4.8 | Laptop computer | Pen | Stapler |
| **Farm** | 1.9 | 5.9 | Tractor | Combine | Fork |
| **Street** | 1.5 | 6.0 | Car | Road sign | Traffic lights |
| **Bathroom** | 1.6 | 7.3 | Towel | Soap | Sink |
| **Kitchen** | 2.3 | 6.9 | Pot | Oven | Refrigerator |
| **Supermarket** | 1.1 | 3.9 | Shopping cart | Shopping basket | Cash register |
| **Garage** | 1.1 | 3.6 | Damaged car | Adjustable wrench | Tire |
| **Classroom** | 3.4 | 6.1 | Black board | Chalks | Pencil case |
| **Church** | 3.4 | 5.5 | Church bench | Crucifix | Altar |
| **Construction site** | 0.7 | 5.2 | Site helmet | Crane | Shovel |
| **Airport** | 2.8 | 6.2 | Luggage trolley | Luggage | Airport bench |
| **Plane*** | 4.8 | 6.8 | Plane seats | Airport window | Airplane reactor |
| **Café/Bar** | 3.7 | 7.6 | Beer glass | Coffee cup | Bar table |
| **Tennis court** | 2.0 | 5.8 | Tennis racket | Tennis ball | Tennis net |
| **Swimming pool*** | 4.5 | 7.2 | Swimsuit | Diving board | Pool ladder |

*Contexts for which the intensity measure does not correspond exactly to their binary classification; these contexts were removed in the additional control analysis.

**Figure S1**

Results of the fMRI analyses repeated while removing the four contexts which were misclassified according purely to the *a posteriori* validation study.

**a)** Brain regions where there is a significant interaction between affective and predictive values ($p < .05$, two-tailed, FWER-corrected at the cluster level with a primary threshold of $p < .001$). These brain regions are: left and right cunei.

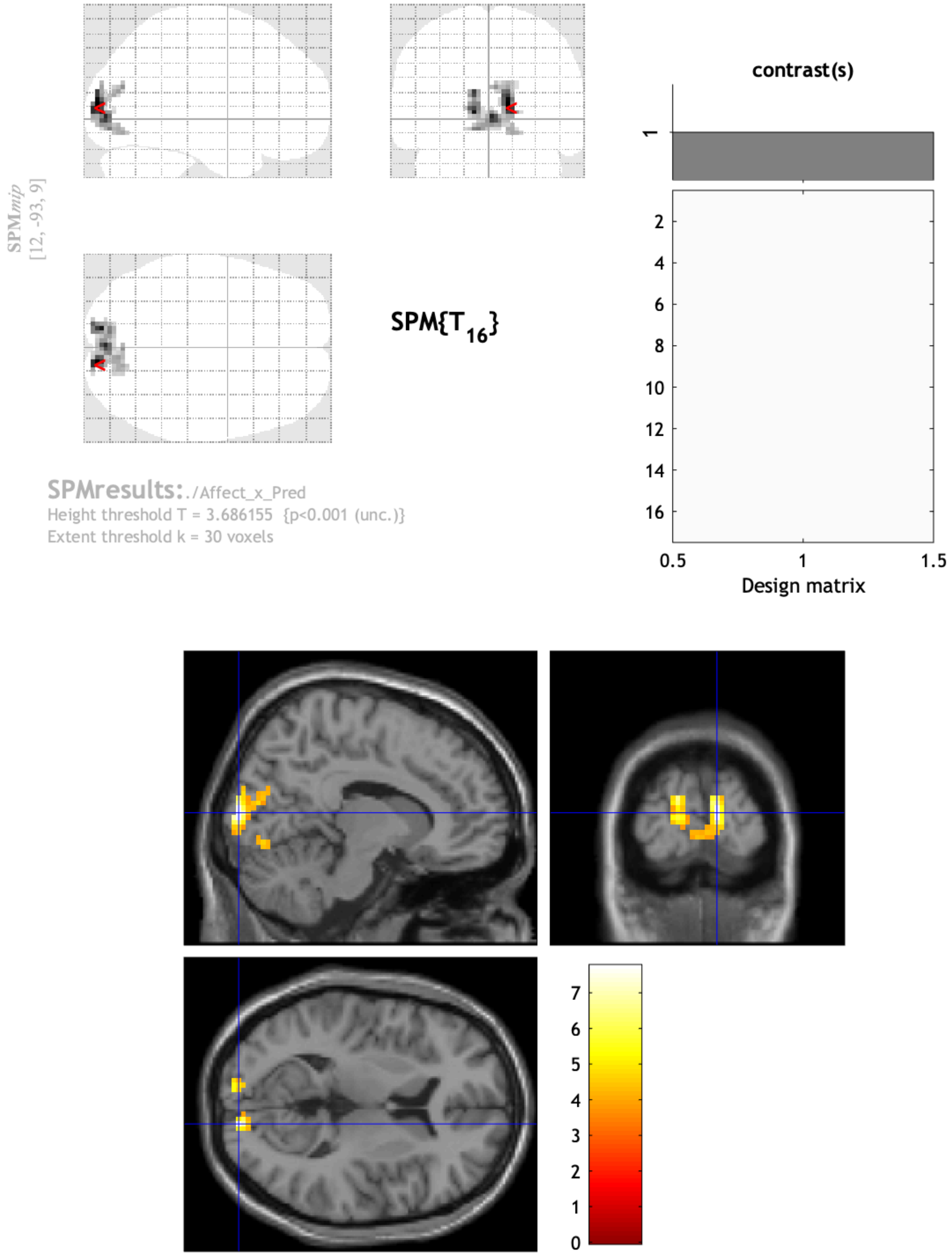

**b)** Brain regions significantly more activated ($p < .05$, two-tailed, FWER-corrected at the cluster level with a primary threshold of $p < .001$) when objects preceded by expectations (Pred+noPred) are viewed than when objects alone (noCont) are viewed. These brain regions are: left precuneus, right precuneus, left middle occipital gyrus, left middle temporal gyrus, left angular gyrus and right angular gyrus. With the exception of the left middle and inferior frontal gyri, these regions are the same as in the main analysis.

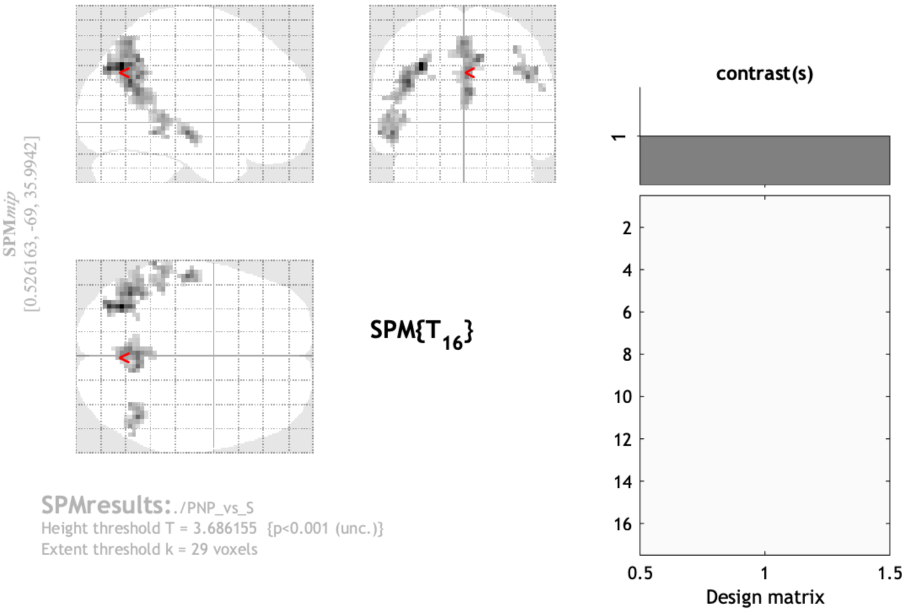


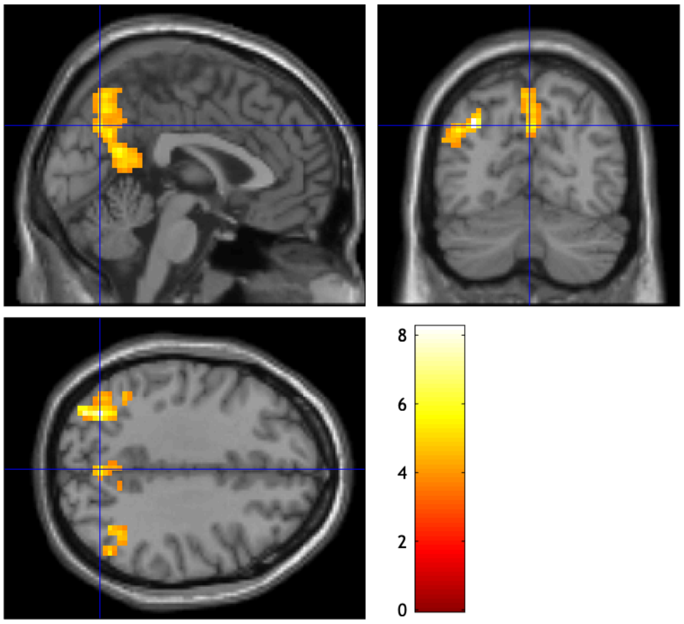

**c)** Brain regions significantly more activated ($p < .05$, two-tailed, FWER-corrected at the cluster level with a primary threshold of $p < .001$) when objects alone (noCont) are viewed than when objects preceded by expectations (Pred+noPred) are. These brain regions are: left middle occipital gyrus, right middle occipital gyrus and right superior occipital gyrus. These regions are qualitatively the same as in the main analysis.

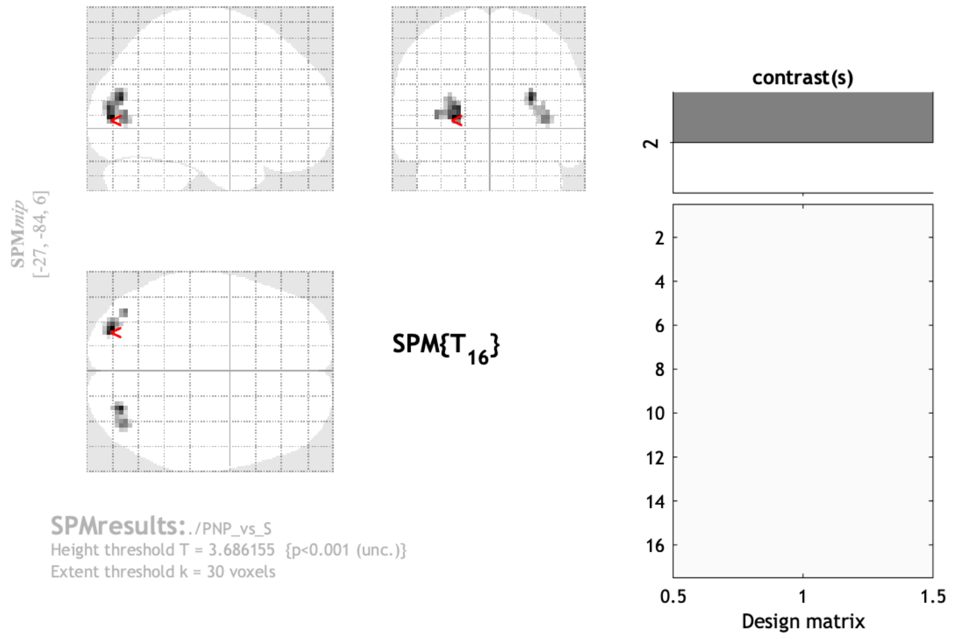


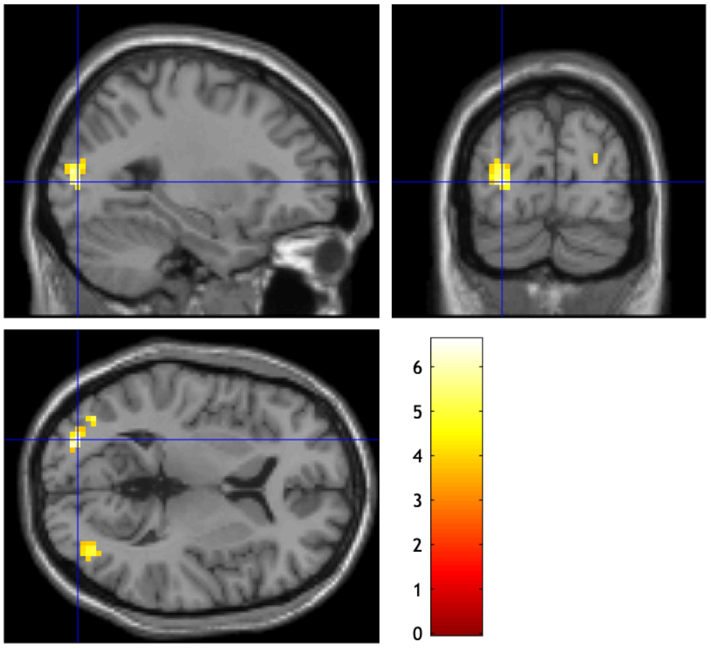

**Figure S2**

Brain regions significantly more activated (*p* < .05, two-tailed, FWER-corrected at the cluster level with a primary threshold of *p* < .001) when regular scene images are viewed than when phase-scrambled scenes are. These regions are: right parahippocampal gyrus, right lingual gyrus, left fusiform gyrus, left lingual gyrus, precuneus, right superior parietal gyrus. These regions are only partly overlapping with the ones uncovered when contrasting predicted and non-predicted objects to objects preceded by a scrambled image.

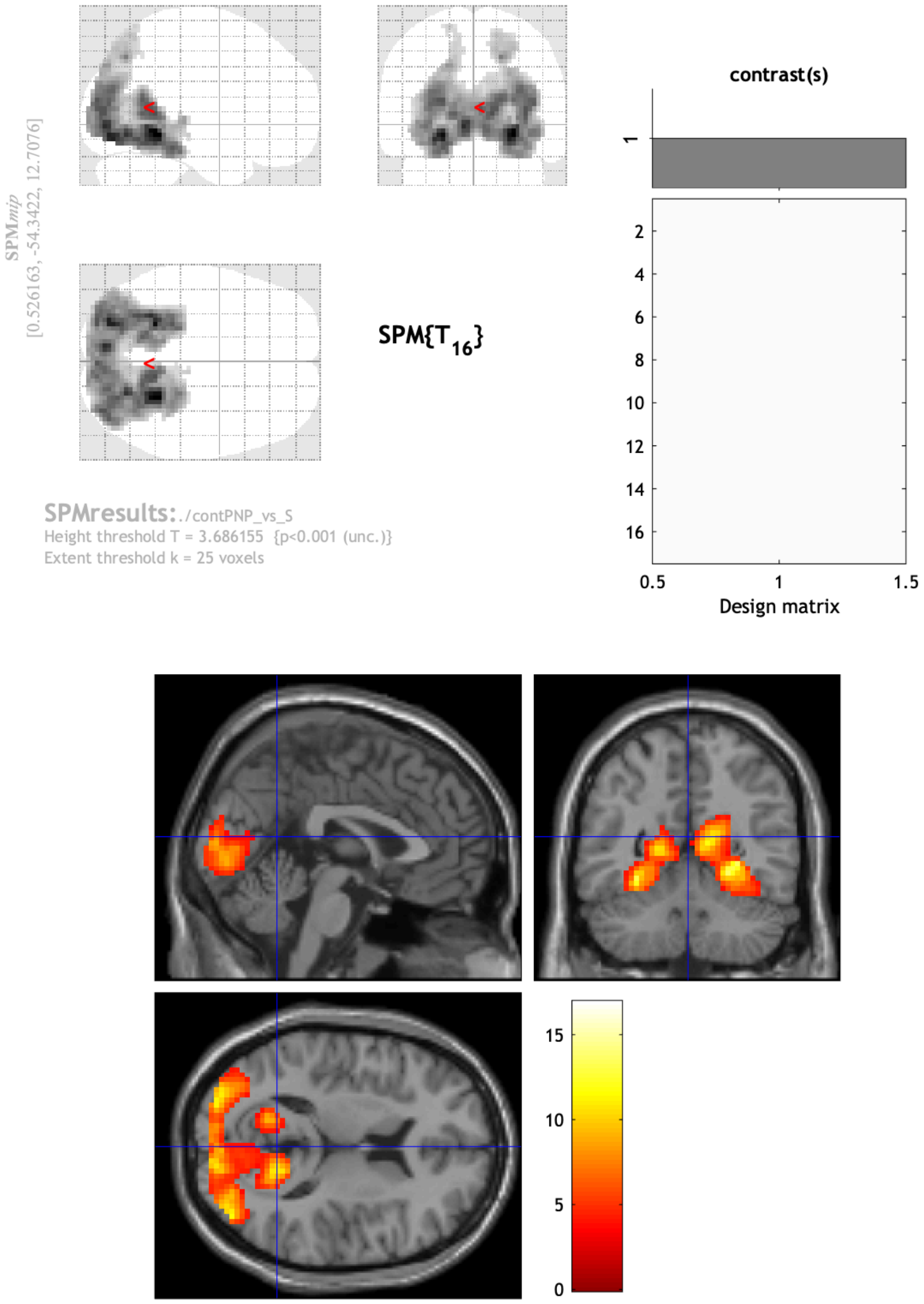

**Figure S3**

Brain region significantly more activated ($p < .05$, two-tailed, FWER-corrected at the cluster level with a primary threshold of $p < .001$) when affective scenes are viewed than when neutral scenes are. This region is the right inferior temporal gyrus.

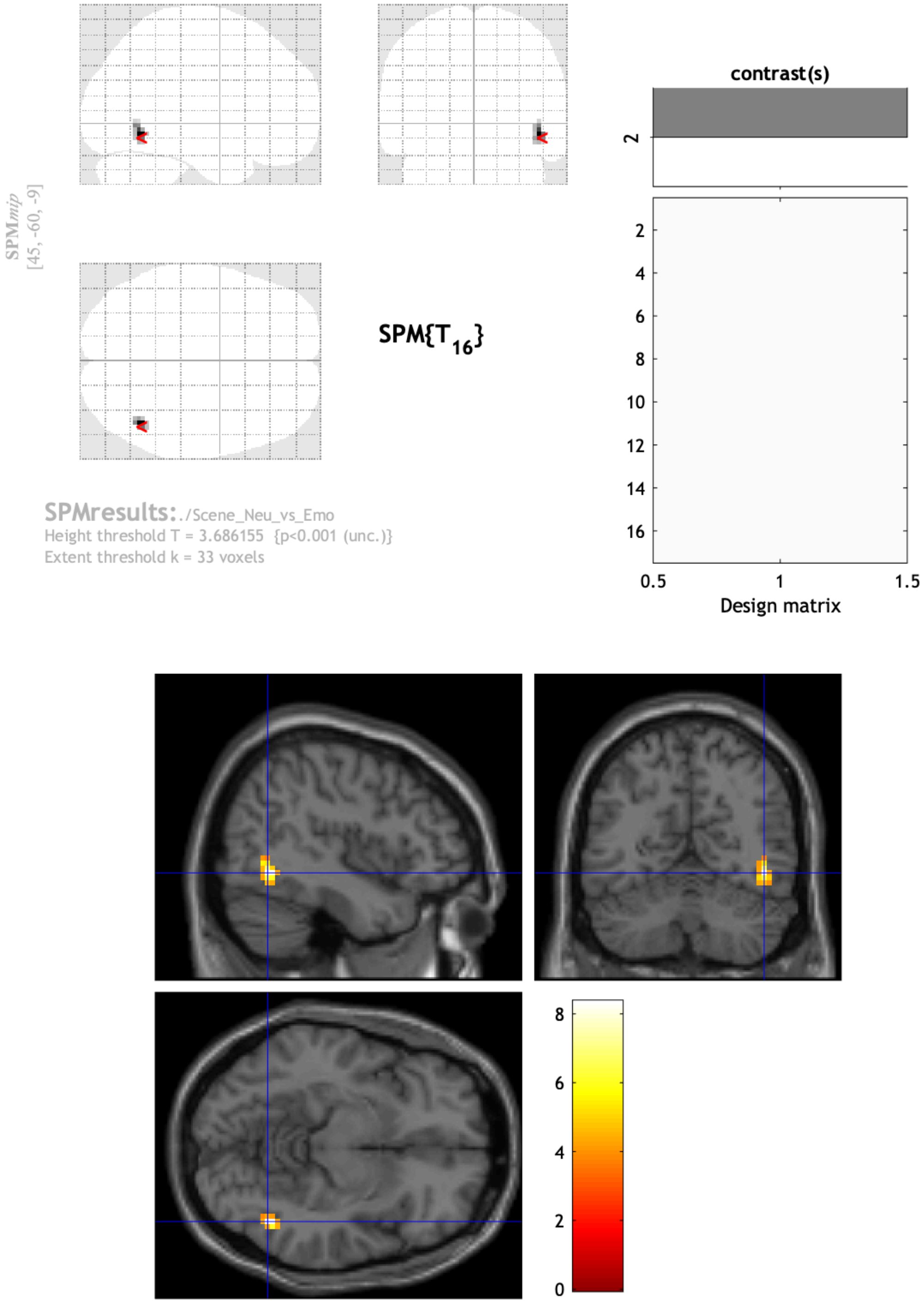

**Figure S4**

Brain regions significantly activated ($p < .05$, two-tailed, FWER-corrected at the cluster level with a primary threshold of $p < .001$) when contrasting all objects together (Pred+noPred+noCont) to the baseline. These regions are: left fusiform gyrus, right fusiform gyrus, left inferior occipital gyrus, left frontal inferior gyrus (operculum part), the left middle frontal gyrus, the precuneus, the left superior parietal gyrus and the left superior frontal gyrus. These regions include part of the lateral occipital complex (posterior fusiform gyri and lateral occipital cortices), typically activated when objects are perceived (Grill-Spector et al., 2001).

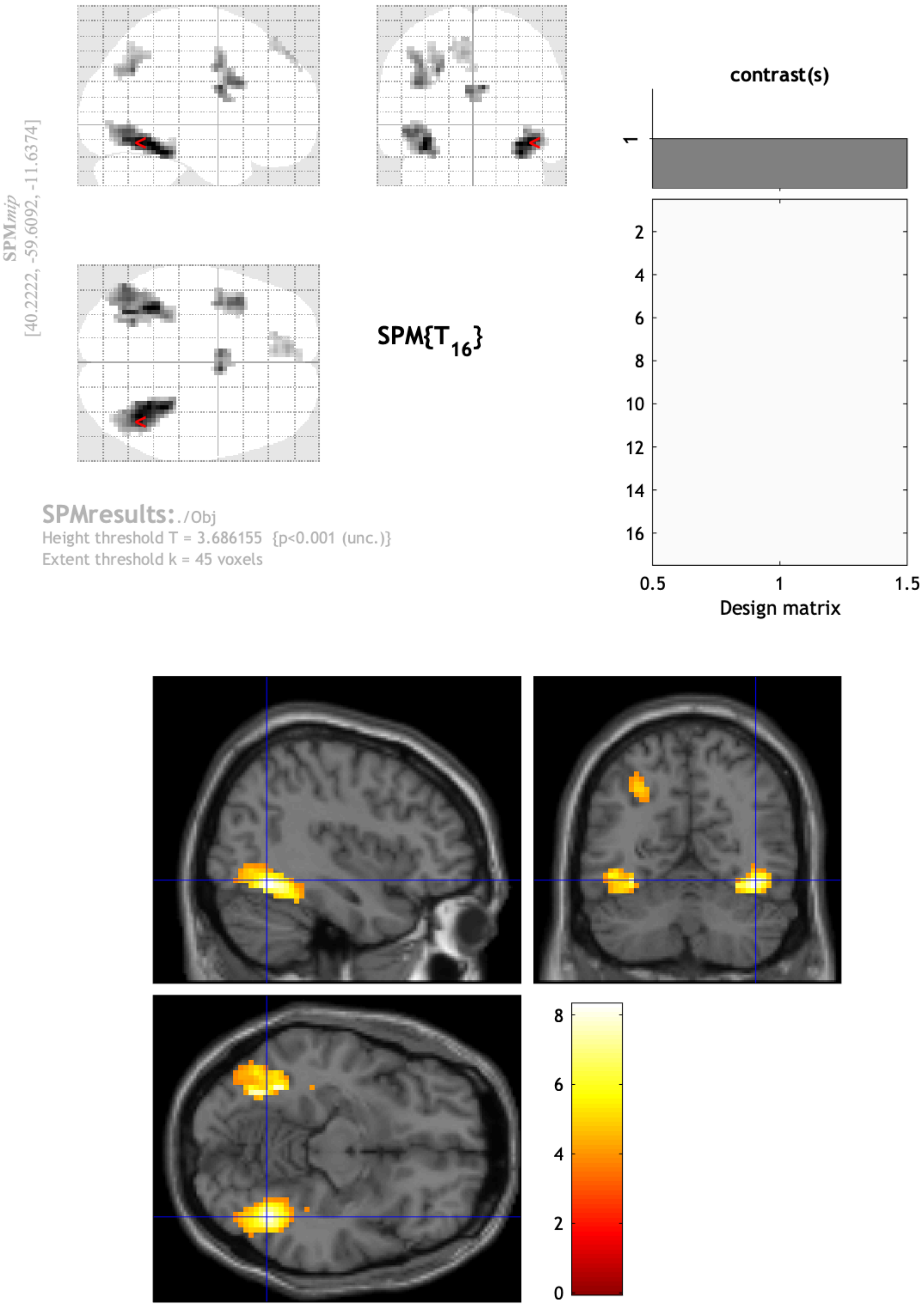
